# Phylogenetic profiling suggests early origin of the core subunits of Polycomb Repressive Complex 2 (PRC2)

**DOI:** 10.1101/2021.07.16.452543

**Authors:** Abdoallah Sharaf, Mallika Vijayanathan, Miroslav Oborník, Iva Mozgová

## Abstract

Polycomb Repressive Complex 2 (PRC2) is involved in establishing transcriptionally silent chromatin states through its ability to methylate lysine 27 of histone H3 by the catalytic subunit Enhancer of zeste [E(z)]. Polycomb group (PcG) proteins play a crucial role in the maintenance of cell identity and in developmental regulation. Previously, the diversity of PRC2 subunits within some eukaryotic lineages has been reported and its presence in early eukaryotic evolution has been hypothesized. So far however, systematic survey of the presence of PRC2 subunits in species of all eukaryotic lineages is missing. Here, we report the diversity of PRC2 core subunit proteins in different eukaryotic supergroups with emphasis on the early-diverged lineages and explore the molecular evolution of PRC2 subunits by phylogenetics. In detail, we investigate the SET-domain protein sequences and their evolution across the four domains of life and particularly focus on the structural diversity of the SET-domain subfamily containing E(z), the catalytic subunit of PRC2. We show that PRC2 subunits are already present in early eukaryotic lineages, strengthening the support for PRC2 emergence prior to diversification of eukaryotes. We identify a common presence of E(z) and ESC, suggesting that Su(z)12 may have emerged later and/or may be dispensable from the evolutionarily conserved functional core of PRC2. Furthermore, our results broaden our understanding of the E(z) evolution within the SET-domain protein family, suggesting possibilities of function evolution. Through this, we shed light on a possible emerging point of the PRC2 and the evolution of its function in eukaryotes.

## Introduction

Polycomb-group (PcG) proteins are evolutionarily conserved chromatin-associated multisubunit complexes that are involved in the regulation of key developmental programs in multicellular organisms. PcG proteins were first discovered as regulators of homeotic (HOX) gene transcription and developmental body patterning in *Drosophila melanogaster* (Lewis 1978; Struhl 1981; Schuettengruber et al. 2017). PcG proteins serve to selectively repress gene transcription by forming multi-subunit complexes called “Polycomb Repressive Complexes (PRC)”, which are sub-categorized in to three major groups: PRC1, PRC2 and Polycomb Repressive DeUBiquitinase “PR-DUB” (Chittock et al. 2017). Among these PRCs, PRC2 is the best-well studied complex that catalyzes repressive methylation of histone H3 on lysine 27 (H3K27) (Cao et al. 2002; Kuzmichev et al. 2002). On multiple genomic sites, PRC2 acts in concert with the H2A E3 ubiquitin-ligase complex PRC1 by recognizing and/or promoting PRC1-deposited histone mark H2AK119ub1 to mediate relatively stable transcriptional repression associated with chromatin compaction (Simon and Kingston 2009; Margueron and Reinberg 2011; Kasinath et al. 2021). By targeting different gene sets in time and space, PRC2 represents a major evolutionarily conserved epigenetic repressive system that governs cell identity and development in multicellular eukaryotes (Mozgova et al. 2015; Yu et al. 2019; Yan et al. 2020; Piunti and Shilatifard 2021).

The core of⍰PRC2 comprises four subunits, all of which are highly conserved in higher eukaryotes (Margueron and Reinberg 2011; Bauer et al. 2016; Bratkowski et al. 2018). Interestingly, higher eukaryotes make use of multiple alternative versions of some of the subunits, which can confer diverse properties and/or functions to the complex (Margueron and Reinberg 2011). The fact that PRC2 components are well conserved in multicellular eukaryotes and absent in unicellular yeast models (*Saccharomyces cerevisiae* and *Schizosaccharomyces pombe*) initially indicated that PRC2 may have co-evolved with multicellularity (Trojer and Reinberg 2006; Köhler and Villar 2008; Veerappan et al. 2008). This notion has since been challenged by the identification of PRC2 subunits and H3K27 methylation in several unicellular eukaryotes spanning different eukaryotic supergroups (Shaver et al. 2010; Dumesic et al. 2015; Veluchamy et al. 2015; Mikulski et al. 2017; Zhao et al. 2020). In unicellular species, PRC2 has been functionally connected to the repression of transposable elements (Shaver et al. 2010; Frapporti et al. 2019) and the determination of cell identity (Zhao et al. 2021).

Based on these findings, PcG proteins are now hypothesized to have been present in the Last Eukaryote Common Ancestor (LECA) and PRC2 to have emerged through divergent evolution, but to have been secondarily lost in the model yeasts (Shaver et al. 2010; Zhao et al. 2020). To date, the diversity of PRC2-core subunits has only been studied in selected eukaryotic lineages, leaving open the question of evolutionary relationships between the homologs of each subunit and the point of PRC2 emergence.

In *D. melanogaster*, PRC2 core is composed of four protein subunits: the catalytic subunit Enhancer of zeste [E(z)] - a histone methyltransferase (HMT) responsible for repressive H3K27 methylation, the Suppressor of zeste 12 [Su(z)12] - a C2H2-type zinc finger protein, and two different WD40 repeat (WDR) domain proteins, namely Extra sex combs (ESC) and Nucleosome remodeling factor (Nurf55) (Birve et al. 2001; Kuzmichev et al. 2002; Nekrasov et al. 2005; Tie et al. 2007; Müller and Verrijzer 2009). The subunit homologs in human are EZH1/EZH2, SUZ12, embryonic ectoderm development (EED) and RBBP4/7 (retinoblastoma binding protein) (Chammas et al. 2020), respectively. Plant PRC2 evolved towards a higher number of homologs, with three E(z), three Su(z)12, one ESC and up to 5 Nurf55 subunit homologs in *Arabidopsis thaliana* (Hennig and Derkacheva 2009). In metazoans (multicellular animals), PRC2 also acts as a H3K27 mono- and di-methyltransferase, and H3K27 tri-methyltransferase activity of the PRC2 can be enhanced by the presence of extra subunits such as Polycomb-like (Pcl) (Nekrasov et al. 2007; Cao et al. 2008; Sarma et al. 2008; Müller and Verrijzer 2009). In plants, however, PRC2 mediates H3K27 tri-methylation while mono-methylation is carried out by the ARABIDOPSIS TRITHORAX-RELATED proteins ATXR5 and ATXR6 (Jacob et al. 2009), that act separately from the PRC2. H3K27me3, the conserved hallmark of PRC2 activity, is generally connected to transcriptional repression (Wiles and Selker 2017), occupying genes located in facultative heterochromatin (Lee et al. 2006; Schwartz et al. 2006; Zhang et al. 2007; Liu et al. 2011) but also, repetitive sequences in constitutive heterochromatin of some species (Jamieson et al. 2013; Veluchamy et al. 2015; Mikulski et al. 2017; Montgomery et al. 2020).

The catalytic activity of the E(z) subunit is conferred by its structurally conserved SET (Su(var)3-9, Enhancer-of-zeste and Trithorax) domain (Dillon et al. 2005; Zhang and Ma 2012; Batra et al. 2020) of 120-130 amino acids in length, which was first identified in several chromatin-associated proteins (Jones and Gelbart 1993; Tschiersch et al. 1994; Jenuwein et al. 1998). The SET-domain structure contains β-strands, several α-helices, turns and numerous loops. The N-terminal region is specifically organized as an antiparallel β conformation (Wilson et al. 2002; Yeates 2002). Resembling the three-dimensional structures of other SET-domains, the E(z)-SET domain also possesses a similar structural fold and conformation with a “thread-the-needle” arrangement, called a pseudoknot (Jacobs et al. 2002; Dillon et al. 2005). SET-domain-containing HMT’s have been organized into seven subfamilies characterized by their phylogenetic relationships: E(z), Trx, Ash, SETD, SMYD, PRDM and Suv (Dillon et al. 2005; Ng et al. 2007; Zhang and Ma 2012; Vervoort et al. 2016; Sarma and Lodha 2017; Zhou et al. 2020). The Suv and E(z) subfamilies are involved in gene repression, the Trx, SETD, and Ash subfamilies are positive regulators, and the members of SMYD and PRDM subfamilies mediate both gene repression and activation (Dillon et al. 2005; Hohenauer and Moore 2012; Zhang and Ma 2012; Carvalho et al. 2013; Vervoort et al. 2016; Tracy et al. 2018).

SET-domain proteins were also identified in bacteria and archaea. Initially, these were described as a paradigm for Horizontal Gene Transfer (HGT) from eukaryote hosts to symbiotic or pathogenic prokaryotes (Stephens et al. 1998; Aravind and Iyer 2003). The availability of sequenced bacterial genomes however uncovered the presence of SET-domain proteins also in free-living and environmental species (Alvarez-Venegas et al. 2007; Alvarez-Venegas 2014), indicating ancient evolutionary origin of SET-domain proteins. SET-domain proteins of bacterial pathogens can activate several eukaryotic cell functions (Alvarez-Venegas et al. 2007; Murata et al. 2007; Li et al. 2013; Alvarez-Venegas 2014). Recent evidence also supports a eukaryotic chromatin-related function of some bacterial SET-domain proteins (Aravind et al. 2011; Alvarez-Venegas 2014). Archaeal SET-domain proteins selectively methylate the archaeal histone H4 at Lys37, suggesting the existence of chromatin methylation before the separation of the archaeal and eukaryotic domains (Manzur and Zhou 2005; Alvarez-Venegas 2014). Yet, the presence of SET-domain proteins in the Asgard (or Asgardarchaeota) group, a separate domain of life representing the closest prokaryotic relatives of eukaryotes (Eme et al. 2017), has not been determined.

Here, using a newly developed automated computational pipeline “PcG-finder”, we identify and report the divergent paralogues and reconstruct the phylogeny of PRC2-core subunits in all eukaryotic lineages, including several unicellular organisms hypothesized to be proximal to the root of eukaryotic supergroups. We also determine a likely evolutionary emergence point for each of the PRC2 core subunits. For the first time we investigate the SET-domain protein family evolution within the four domains of life (Asgard, Archaea, Bacteria, and Eukaryote), with an emphasis on the structure-function relationship of the E(z) homolog SET domains. Identifying E(z) homologs in early diverged groups, our findings provide significant support for PRC2 emergence prior to the diversification of eukaryotic lineages.

## Results and Discussion

### Phylogenetic distribution of putative PRC2 subunit homologs

PcG protein complexes in non-model and unicellular species are less thoroughly investigated than in multicellular model animals or plants. PRC2 components were first discovered in microalgae in 2010 (Shaver et al., 2010). Most recently, the diversity of Polycomb and Trithorax complexes was investigated in single-celled species represented in the Marine Microbial Eukaryote Transcriptome Sequencing Project (MMETSP) database (Keeling et al. 2014; Zhao et al. 2020), where major eukaryotic lineages are represented but several groups, including Metamonada, Discoba (Eutetramitia), Opisthokonta (Metazoa), Viridiplantae (Embryophyta), Rhodelphidia, Colponemidia (Alveolata) and Eustigmatophyceae or Oomycota (Stramenopila) were missing or not studied. In quest for the origin of PRC2 subunits in eukaryotes, we used publicly available genomic and transcriptomic data from species within all eukaryotic supergroups including Discoba and Metamonada, groups hypothetically proximal to the root (Adl et al. 2012; Burki et al. 2020). We used the full-length reference protein sequences of PRC2 subunits from *D. melanogaster* as a query and retained only those sequences that contained catalytic or conserved domains with similarity to the respective reference sequence. In total, genomes or transcriptomes of 283 species were explored here (fig. 1; table S1A, , Supplementary Material online), leading to the identification of predicted protein isoforms of the four core PRC2 subunit homologs. Homologs of at least one of the PRC2 subunits were identified in 262 (92.6%) of these species, and all the subunit homologs were found in 56 (19.8%) species (fig. 1; table S1B, Supplementary Material online).

**Figure 1.**
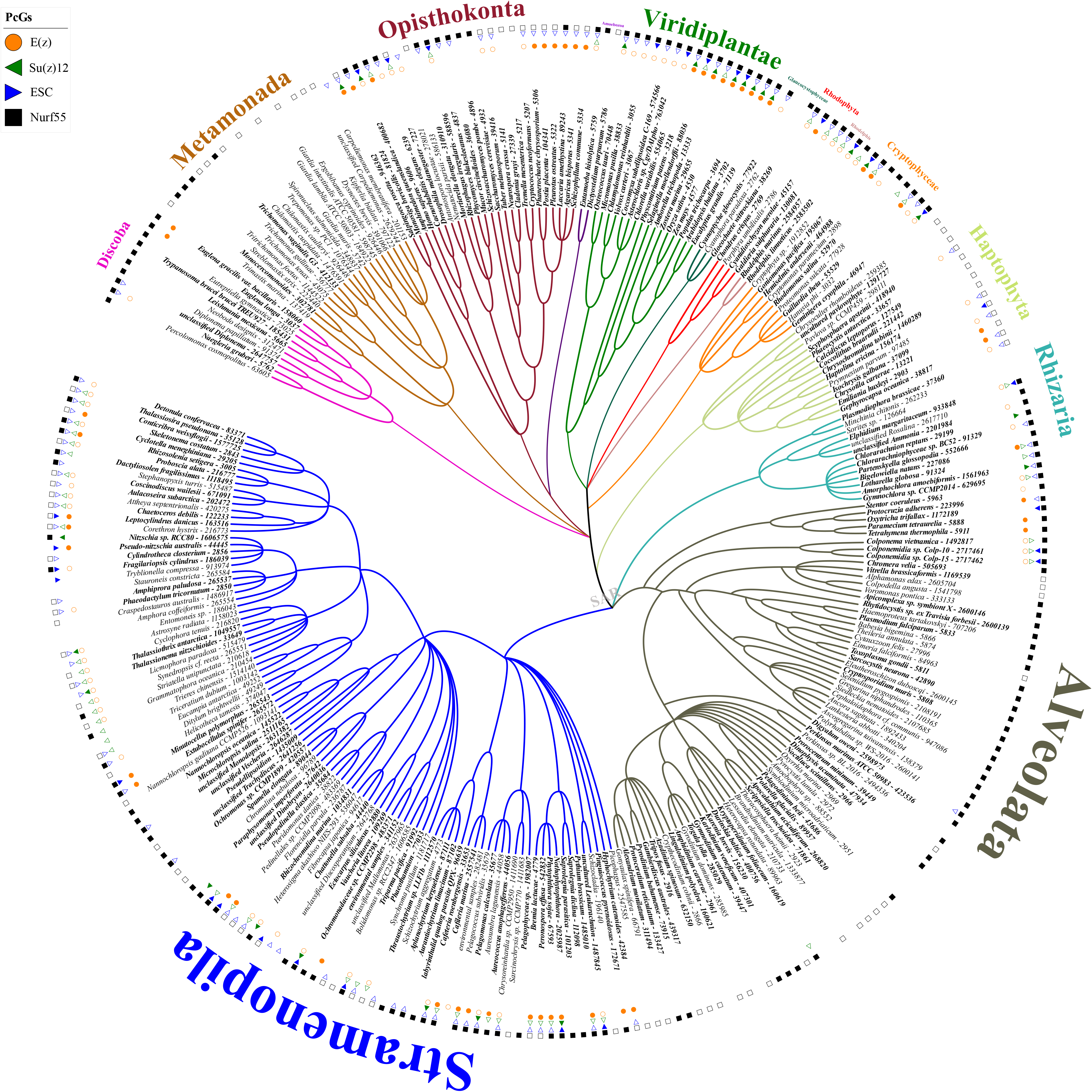
Species cladogram shows the distribution of PRC2 subunit homologs across eukaryotic lineages. Species with BUSCO score C ≥ 50 are marked in bold. Empty symbols indicate one identified homolog, filled symbols indicate multiple identified homologs. Homologs may originate form one or more genomic loci.

### E(z) subunit

E(z) is the catalytic H3K27 methyltransferase subunit of PRC2 (Goodrich et al. 1997; Cao et al. 2002; Czermin et al. 2002; Kuzmichev et al. 2002; Müller et al. 2002). One homolog is found in *D. melanogaster* (E(z)), two in mammals (EZH1 and EZH2) and three in *Arabidopsis thaliana*, (CURLY LEAF (CLF) (Goodrich et al. 1997), SWINGER (SWN) (Luo et al. 1999; Chanvivattana et al. 2004) and MEDEA (MEA) (Grossniklaus et al. 1998). The E(z) subunit is found in representatives of major eukaryotic supergroups, and it is highly conserved in stramenopiles and chlorophyte microalgae (Zhao et al. 2020).

Out of 283 species studied, E(z) was identified in 119 (42%) species. In agreement with previous studies, E(z) was identified in all studied representatives of the green lineage (Viridiplantae) (Shaver et al. 2010; Huang et al. 2017; Ni et al. 2019) (fig. 1; table S1C, Supplementary Material online). The E(z) was differently identified or not in representatives of all other eukaryotic lineages, but it was absent from all studied species of Metamonada and Amoebozoa. Interestingly, E(z) homologs were identified only in euglenoids and the heterolobosean free-living amoeboflagellate *Naegleria gruberi* within the Discoba lineage. Among the alveolates, E(z) homologs were identified only in the ciliates and in the recently discovered family colponemidia (Tikhonenkov et al. 2020) (fig. 1; table S1C, Supplementary Material online). The number of E(z) homolog isoforms ranged from 1 to 19 found in the plant *Populus trichocarpa*. As our study is based on inferred protein sequences from transcripts, protein isoforms are identified that may represent alternative transcripts of a single gene locus. Nevertheless, SET-domain protein expansion was observed and attributed to whole-genome duplication events in *P. trichocarpa* (Lei et al. 2012), offering possible explanation for a large number of homologs. In contrast to Huang and colleagues (Huang et al. 2017), we identified only one E(z) homolog in the studied chlorophytes, and two or more homolog isoforms in all embryophytes (fig. 1; table S1C, Supplementary Material online).

Earlier, absence of PRC2 subunits in some eukaryotic lineages was observed (Zhao et al. 2020) and was attributed to low or undetected expression in the transcriptomic data from the MMETSP database (Keeling et al. 2014). This is not likely to apply here for at least some species because in 55% and 28% of the species of metamonads and alveolates respectively, the genomic data used showed very high coverage (500x) (table S1A, Supplementary Material online). In order to further support the potential absence of E(z) in some species compared to others where E(z) was found, we quantitatively evaluated the starting datasets for completeness in terms of expected gene content using BUSCO software (https://busco.ezlab.org) (Seppey et al. 2019). The BUSCO scores show that 182 (64.3%) of the 283 used datasets show more than 50% completeness (fig. 1 – species in bold; table S1A, Supplementary Material online). In alveolates, a higher proportion of species with E(z) identified would be expected if E(z) was generally present, suggesting that E(z) may indeed be missing in a considerable subset of species within this group. Nevertheless, we cannot rule out the possibility of insufficient data quality or poor structural annotation of these datasets that would impede the identification of homologs in the individual species. As an obvious example, the previously identified E(z) homolog (Shaver et al. 2010; Huang et al. 2017; Mikulski et al. 2017) based on an older *Chlamydomonas reinhardtii* genome assembly (v4) (Merchant et al. 2007), is a short sequence (56 aa) that lacks the catalytic SET-domain. However, we could identify a longer E(z) homolog sequence of 793 aa using the new version of the *C. reinhardtii* genome (v5.6) (O’Donnell et al. 2020). True absence of E(z) in species or whole lineages will need to be further confirmed also experimentally and it will be interesting to elucidate alternative pathways that may fulfill the function of H3K27me3-mediated repression or identify alternative SET-domain proteins that may take the function of E(z). Importantly, we show that E(z) homologs are found in one of the lineages closest to the eukaryotic root (Discoba) (Adl et al. 2012; Burki et al. 2020): This provides significant support for the hypothesis of E(z) presence in the LECA and emergence preceding the diversification of eukaryotes (Shaver et al. 2010; Zhao et al. 2020). In addition, loss of E(z) is a recurring event in the eukaryotic evolution that seems to largely affect the alveolates. The loss of E(z) in alveolates may have occurred following the emergence of colponemids (fig. 1; table S1C, Supplementary Material online).

We next computed a eukaryotic-E(z) phylogenetic ML tree based on 814 sequences comprising sequences identified using the PcG finder and sequences previously identified in KOG1079. The unrooted phylogenetic tree clustered E(z) homologs into five clades (Clade I – V, fig. 2A; table S1E, Supplementary Material online). Clade I contains mainly fungal sequences; including the E(z) homolog of *Cryptococcus neoformans*, the first budding yeast with identified and characterized PRC2 complex (Dumesic et al. 2015). All euglenoid sequences are grouped in the root of the same clade (fig. 2A). Almost all the ciliate sequences are grouped in the crown of clade II; these sequences were previously identified as a different sub-class of E(z) named the Enhancer-of-zeste-like protein Ezl (Frapporti et al. 2019). The one *Naegleria gruberi* (Discoba) E(z) homolog and all *Cafileria marina* (Jirsová et al. 2019) (stramenopiles) homologs cluster in the same clade. Clade III contains all the rhodophyte sequences, including the recently identified homologs in *Cyanidioschyzon merolae* (Mikulski et al. 2017) and the E(z) homologs of the newly discovered family Rhodelphidae (Gawryluk et al. 2019; Burki et al. 2020) (fig. 2A). The phylogenetic tree from the previous study showed a separation between C. merolae E(z) homolog and other studied E(z) of Viridiplantae and Opisthokonts (Mikulski et al. 2017). This is in line with the observed separation of these homologs in different clades. Interestingly, all stramenopile sequences clustered in the last two clades IV and V. Clade IV contains about half of the stramenopile and all colponemid E(z) homologs, except *Cafileria marina* homologs which cluster in clade II. The second half of the stramenopile sequences cluster near the root of clade V (fig. 2A). Clade V further contains almost all opisthokont sequences with the Eumetazoa homologs (EZH1 and EZH2) in the crown of the same clade. All Cryptophyta, Viridiplantae, Rhizaria and Haptophyta sequences cluster in the root of clade V. We did not identify overlap of species between these five clades, with the exception of 19 stramenopile species that formed an overlap of clades IV and V (Oomycota, Ochrophyta, and Hyphochytriomycota) (fig. S1A; table S2A, Supplementary Material online). This finding supports divergent evolution of E(z) subunit homologs of a single clade within species and/or even whole eukaryotic lineages, perhaps in connection with gene duplication during evolution, as was recognized before within green lineage (Huang et al. 2017; Ni et al. 2019). The divergence of the homologs of the 19 species from the stramenopile groups remains unclear but indicate the co-existence of two separate clades of E(z) homologs in these species, the significance of which remains unclear so far. In summary, the phylogenetic tree of eukaryotic E(z) homologs highlights the existence of five different sub-classes of E(z) sequences, which are partially lineage-specific.

**Figure 2.**
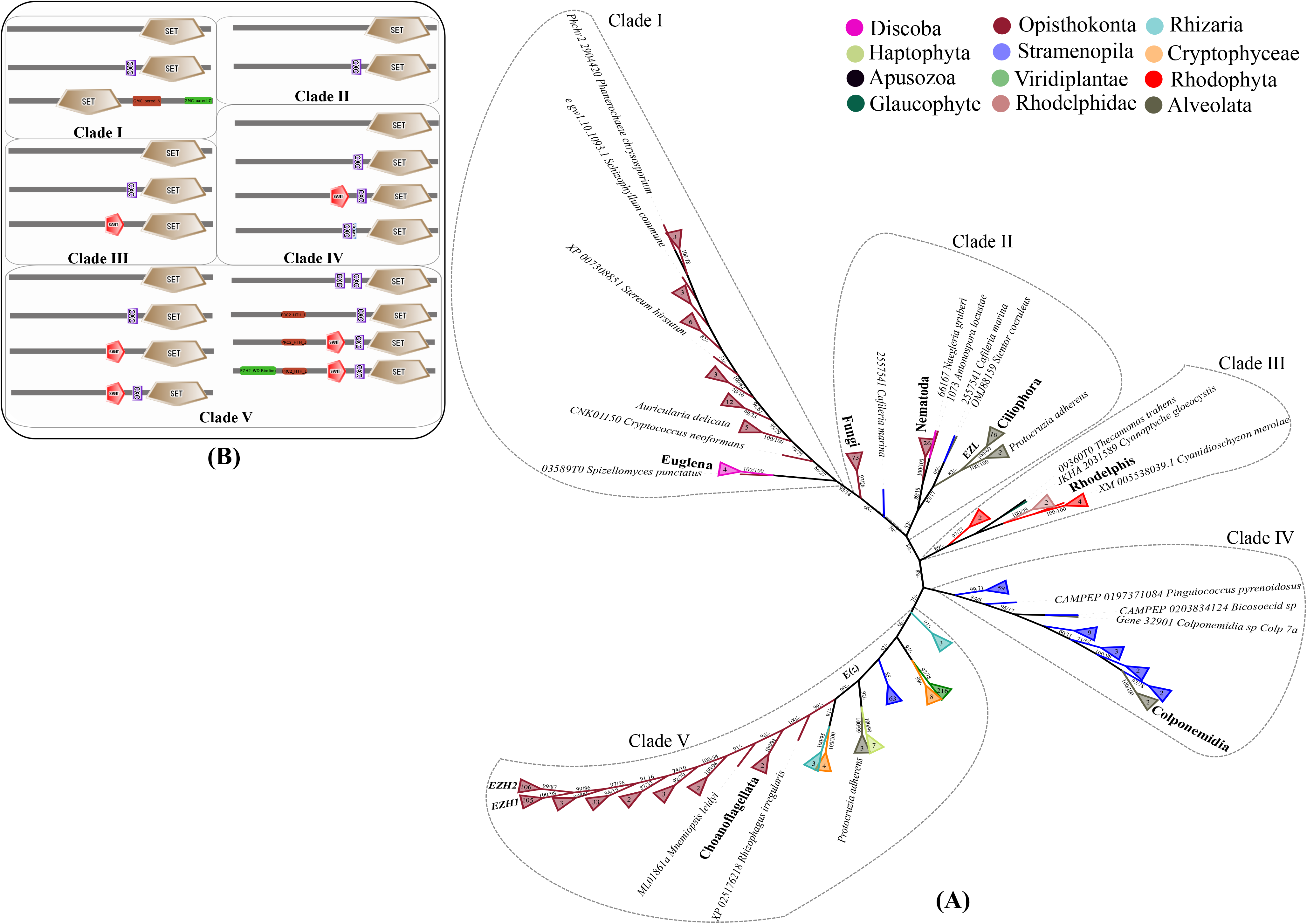
A ML-eukaryotic phylogenetic tree of E(z) homologs showing the evolutionary relationship and domain architectures. **A)** The tree was constructed using the whole protein sequence and the maximum likelihood branch support values are given in % (IQ-TREE/RAxML-NG). **B)** The domain architectures of the full-length proteins were drawn based on the sequences representatives of each clade and a search against SMART and Pfam databases.

Previous domain organization and phylogenetic relationship analyses suggested that E(z) proteins may be involved in H3K27me3 deposition (Huang et al. 2017). Having identified five clades of the E(z) homologs based on the full-length protein sequences, we were interested whether the domain architecture differentiates homologs in respective clades. From each of the five clades of E(z) homologs, we selected sequences that best represent the full diversity of the multiple sequences alignments (MSA) using hhfilter script from HH-suite3 software (https://github.com/soedinglab/hh-suite) (Steinegger et al. 2019) (fig. 2B; S2B, Supplementary Material online). In general, we identified three well-conserved domains that are associated with different E(z) homologs. The SET (PF00856) domain is a conserved sequence of 130–150 amino acids that serves as the catalytic domain of lysine methyltransferases (Qian and Zhou 2006; Huang et al. 2011). The SANT (SM00717) domain is found in several other chromatin remodeling proteins with various functions, such as DNA-binding, histone tail-binding, and protein-protein interactions (Zhang et al. 2006). CXC (PF03638) is an ~65-residue cysteine-rich region preceding the SET domain and forms a pre-SET subdomain (Ketel et al. 2005). Further, the CXC domain interacts with nucleosomal DNA (Robert et al. 1996; Tokodai and Yakushiji 2019).

Domain architecture screening shows that representative sequences of E(z) homologs from the five clades contain different combinations of the three domains. Clades IV and V contain E(z) homologs with (i) only SET domain, (ii) CXC and SET, or (iii) SANT, CXC and SET domains (fig. 2B, supplementary table S2B). The combination of SANT and SET domains was found in clades III and V. Moreover, proteins of clade I and II contain species closest to the supposed eukaryotic root, suggesting that these may represent the most ancient version of the E(z) protein. These clades contain homologs of *N. gruberi* and all euglenoids (fig. 1 and 2B; table S2B, Supplementary Material online). Clade I and II representative proteins contain either only the SET domain or two sequentially arranged canonical domains (CXC and SET) (fig. 2B; table S2B, Supplementary Material online). One exception is the clade I representative E(z) homolog of the fungus *Auricularia delicata* (160860_XP_007350139_1_KOG_Auricularia_delicata), that contains the N-terminal and C-terminal (GMC_oxred_N “PF00732” and GMC_oxred_C “PF05199”) domains of glucose-methanol-choline oxidoreductase family (GMC oxidoreductase) (Bannwarth et al. 2004). The SANT domain emerged in Clade III N-terminally of the arranged canonical domains (SANT and SET) (fig. 2B; table S2B, Supplementary Material online) and is also found in proteins of clades IV and V. Additionally, the three sequentially arranged canonical domains (SANT, CXC, and SET) can be combined with the nucleic acid-binding Zinc finger (Znf) (ZnF_C2HC “SM00343”) domain (Gamsjaeger et al. 2007) as exemplified by a representative homolog from clade IV. More complex domain architectures were observed within proteins of clade V. This clade contains all higher eukaryote sequences, including eumetazoan homologs (EZH1 and EZH2). The PRC2-specific and DNA-binding domains such as the binding site to EED subunit (EZH2_WD-Binding “PF11616”) (Han et al. 2007) and PRC2 tri-helical domain (PRC2_HTH_1 “PF18118”) were identified (fig. 2B; table S2B, Supplementary Material online). The latter domain makes up part of the N-lobe which is involved in the repressive function of PRC2 (Justin et al. 2016). Protein domains diversity within E(z) has been observed before within the green lineage (Huang et al. 2017; Sarma and Lodha 2017) and other eukaryotic lineages (Shaver et al. 2010; Zhao et al. 2020). In summary, we suggest that the generally present SET domain may be present in combination with the CXC domain in all clades of the E(z) homologs, including the hypothetically most ancient clades I and II. The SANT domain may be present in the E(z) homologs of clades III, IV and V.

### Su(z)12 subunit

Su(z)12 is a Cys2 His2 (C2H2) zinc-finger and VEFS-domain (VRN2/EMF2/FIS2/Su(z)12) core subunit of PRC2 that is conserved in Drosophila, mammals and plants (Birve et al. 2001; Huang et al. 2017). One Su(z)12 homolog is found in either Drosophila or mammals, while three are present in *Arabidopsis thaliana* [EMBRYONIC FLOWER 2 (EMF2), FERTILIZATION-INDEPENDENT SEED 2 (FIS2), and VERNALIZATION 2 (VRN2)], where EMF2 is considered most ancestral (Chen et al. 2009). In animals and plants, Su(z)12 is required for PRC2-mediated developmental regulation and enzymatic activity of PRC2 (Luo et al. 2000; Gendall et al. 2001; Yoshida et al. 2001; Cao and Zhang 2004; Pasini et al. 2004; Ketel et al. 2005; Chen et al. 2008). The Su(z)12 subunit was recently shown to be conserved in the SAR group (Stramenopila, Alveolata, and Rhizaria) as well as chlorophyte microalgae (Zhao et al. 2020).

Out of 283 species representing all eukaryotic lineages, Su(z)12 was identified in 80 (~28.3%) species. In Viridiplantae, the number of Su(z)12 homolog isoforms varied from 1 to a maximum of 13 in the plants *Populus trichocarpa* and the moss *Physcomitrium patens* (table S1D, Supplementary Material online). In *P. patens*, three homologs were identified previously (Chen et al. 2009; Hennig and Derkacheva 2009; Huang et al. 2017). The 13 homologs identified here based on the Phypa V3 genome assembly and annotation of P. patens (Lang et al. 2018) are produced from three loci (LOC112283800, LOC112282854, and LOC112277386), which is in agreement with previous reports. Interestingly, the five core chlorophyte taxa included here lack Su(z)12 homologs (fig. 1; table S1D, Supplementary Material online). Similarly, Su(z)12 absence was previously observed in representatives of core chlorophytes (Hennig and Derkacheva 2009; Shaver et al. 2010; Huang et al. 2017; Zhao et al. 2020). Although the Su(z)12 was identified in most of the eukaryotic lineages, it is not identified in representatives of Discoba, Metamonada, Amoebozoa, Glaucophyta, and Haptophyta (fig. 1; table S1D, Supplementary Material online). In alveolates, Su(z)12 homologs were identified only in colponemids (Tikhonenkov et al. 2020) and the binucleate dinoflagellate *Kryptoperidinium foliaceum*, which contains an endosymbiont nucleus of the diatom origin (Figueroa et al. 2009) (fig. 1; table S1D, Supplementary Material online).

We computed a Su(z)12 Maximum-Likelihood (ML) phylogenetic tree based on 495 sequences identified by PcGs-finder and previously identified Su(z)12 sequences from KOG2350. An unrooted phylogenetic tree successfully separated Su(z)12 homologs of the different eukaryotic lineages (fig. S2A, Supplementary Material online). All the sequences of the green lineage (Viridiplantae) clustered in the crown of the tree while the rhodophyte sequences appeared either between plant and chlorophyte clades (*Galdieria sulphuraria*) or in the proximity of bacterivorus heterokont *Cafileria marina*, thus making Su(z)12 from Rhodophyta and even Archaeplastida polyphyletic (fig. S2A, Supplementary Material online). The only dinoflagellate *K. foliaceum* sequence (CAMPEP_0167906012) clusters with the diatom *Nitzschia sp*. sequences, which complies with the diatom origin of the *K. foliaceum* endosymbiotic nucleus (Figueroa et al. 2009). The unclassified *Colponema* isolate sequences (Colp-10 and Colp-15) cluster together with all cryptophyte sequences (supplementary fig. S2A). *Colponema* is phylogenetically branching in the root of Myzozoa (Dinozoa + Apicomplexa) (Tikhonenkov et al. 2020). Our results suggest the Su(z)12 was lost in Alveolata at the root of Myzozoa.

We next assessed the domain structure of the identified Su(z)12 homologs. The VEFS-Box (PF09733) is found in the C-terminal region of Drosophila Su(z)12. This domain is characterized by an acidic cluster and a tryptophan/methionine-rich sequence domain (Yoshida et al. 2001). In proteins associated with Polycomb repression, the VEFS-Box is associated with a zinc-finger domain located approximately 100 residues towards the N-terminus (Birve et al. 2001). Unlike most C2H2-domain-containing proteins that have multiple C2H2 motifs in tandem (Englbrecht et al. 2004), Su(z)12 contains a single C2H2 motif (Chen et al. 2009). Previously, a conserved sequence of 37 amino acids within the C2H2 domain was found in Su(z)12 homologs from plants and animals (Yoshida et al. 2001). Domain architecture screening shows that most Su(z)12 homologs contain VEFS-Box and the zinc finger domains of Di19-(PF05605) and/or C2H2-type (PF13894) (table S1D, Supplementary Material online). The VEFS-Box domain in Viridiplantae Su(z)12 homologs is present either alone or among two and/or three sequentially arranged canonical domains (VEFS-Box and zinc fingers). VEFS-Box and zinc finger domains are conserved in all other lineages (table S1D, Supplementary Material online). All the identified proteins contain either only the VEFS-Box domain or sequentially arranged canonical domains of zinc fingers and varied domains. These findings that higher-eukaryote homologs have similar domain structures to early eukaryotes indicate that Su(z)12 function is likely conserved across different eukaryotic lineages (table S1D, Supplementary Material online).

New domains such as Tau95 (PF09734) and Kinesin-associated (PF16183) domains emerged in land plants (*P. patens* and *Eucalyptus grandis* respectively) (table S1D, Supplementary Material online). Tau95 is a subunit of TFIIIC1 binding factor, which exerts upstream and downstream control over the TFIIIC-DNA complex by providing the complex with higher stability (Jourdain et al., 2003). The Kinesin-associated domain is a functionally unknown short domain; associated with some kinesin-like proteins. In addition, varied non-conserved domains are observed in the rest of the lineages (table S1D, Supplementary Material online). The relevance of these non-canonical domains for Su(z)12 homolog function remains unknown.

### ESC subunit

ESC is a protein of the WD40 repeat family. Two homologs, ESC and ESC-like, are found in Drosophila (Wang et al. 2006; Ohno et al. 2008), one in mammals (enhanced ectoderm development (EED)) (Cao et al. 2002; Kuzmichev et al. 2002) and one in *Arabidopsis thaliana* (FERTILIZATION INDEPENDENT ENDOSPERM (FIE) (Spillane et al. 2000). Transcripts of ESC were previously not found as widely present in diatoms as transcripts of E(z) or Su(z)12 homologs (Zhao et al. 2020). ESC homologs were previously not found in studied representatives of “Excavata”, several Chromalveolata, Amoebozoa and Opisthokonta (Shaver et al. 2010) but were generally found in species of Archaeplastida (Shaver et al. 2010; Huang et al. 2017). In *Drosophila*, ESC is required for high-affinity binding of the PRC2 to nucleosomes and for H3K27me3 activity in vivo (Nekrasov et al. 2005; Tie et al. 2007). Human EED also interacts with subunits of PRC1, orchestrating the activities of the two complexes (Cao et al. 2014).

ESC was identified in 129 (~45.6%) species of the investigated 283 species. In contrast to all other PRC2 subunits, we identified only one ESC homolog in 101 of the species 129 with ESC homolog (~78.3%) (table S1E, Supplementary Material online). In Metamonada, ESC homologs were identified only in the free-living marine flagellate *Carpediemonas membranifera* (Kolisko et al. 2005). Similarly, it is identified only in the heterolobosean *Naegleria gruberi* and euglenozoan *Diplonema*. Contrary to previous reports (Zhao et al. 2020), we show that ESC homologs are well represented in stramenopiles. Unlike Su(z)12 that was only identified in colponemids within alveolates, ESC homologs were identified in colponemids, ciliates and the most early-branching gregarine *Digyalum oweni* in this family (Janouskovec et al. 2019) (fig. 1; table S1E, Supplementary Material online). These findings suggest that ESC subunit was gradually lost in alveolates being retained only in some ciliates, colponemids and *Digyalum*. The number of ESC homologs varied from one in the mycorrhizal fungus *Rhizophagus irregularis* to five in *Rhodelphis limneticus*, a member of the newly discovered family Rhodelphidae (table S1E, Supplementary Material online) containing heterotrophic flagellated predators, forming a sister group to rhodophytes (Gawryluk et al. 2019; Burki et al. 2020).

Based on the combined datasets from the PcG-finder-identified sequences and previously identified ESC sequences in KOG1034 containing together 535 sequences, we constructed a ML-based phylogenetic tree. In this tree, the ESC homologs clustered into three clades (fig. S2B, Supplementary Material online). Almost all Stramenopila, Rhizaria, and Viridiplantae sequences, as well as all Cryptophyta, Haptophyta, and Rhodophyta sequences, were found in the first clade. The second clade is composed of opisthokonts sequences except for the *Diplonema* homolog in the root and the gregarine *D. oweni* in the crown of the clade (fig. S2B, Supplementary Material online). The third clade mostly contains fungal sequences (fig. S2B, Supplementary Material online). Core chlorophyte sequences grouped together with the Thraustochytrida sequences in the crown of the clade with the *N. gruberi* (Discoba) homolog in the root (fig. S2B, Supplementary Material online). The constructed ESC phylogenetic tree is the most robust ESC tree to date, despite the fact that certain relationships remain unsolved. Similarly, unresolved relationships within microalgae species were reported (Zhao et al. 2020), whereas other studies suggest a monophyletic origin for ESC homologs found in most eukaryotes (Huang et al. 2017).

Similar to what has been observed before (Zhao et al. 2020), we identified up to four WD-40 repeat domains in the ESC proteins (table S1E, Supplementary Material online) that can potentially form a platform for protein-protein interactions. The WD40 domain does not have enzymatic activity (Xu and Min 2011) but forms a β-propeller architecture that binds to the histone H3 tail (Suganuma et al. 2008). Other domains such as Nucleoporin 160 (Nup160) was identified within all lineages except one metamonad homolog (Carpediemonas_membranifera_Gene.9777::gnl) (table S1E, Supplementary Material online). Nup160 or basket nucleoporin plays a critical role in RNA export from the nucleus through the nuclear pore (Vasu et al. 2001). This may support the presence of cytoplasmic PRC2 subunits in Metazoa and plants, suggesting alternative functions of PRC2 subunits (Philipp et al. 2010; Oliva et al. 2016).

### NURF55 subunit

The NURF55 subunit is also an evolutionarily conserved WD40 repeat-containing protein. The number of NURF55 homologs varied from 1 to a maximum of 32 found in the parasitic apicomplexan *Eleutheroschizon duboscqi* (fig. 1; table S1F, Supplementary Material online). NURF55 was identified in 252 (~89%) 283 investigated species. Therefore, NURF55 seems generally present in all studied eukaryotic lineages. Also, out of 252 species with identified NURF55 homologs, 96 (~38.1%) species have one copy of NURF55 (fig. 1; table S1F, Supplementary Material online). The few organisms where NURF55 was not identified may therefore represent false negatives (fig. 1; table S1F, Supplementary Material online). In needs to be considered however, that NURF55 is a part of several different protein complexes, including PRC2, the nucleosome remodeling complex NURF, or the post-replicative chromatin assembly complex CAF1 (Verreault et al. 1996; Martínez-Balbás et al. 1998; Hennig et al. 2005; Köhler and Villar 2008; Wen et al. 2012). Therefore, as in *Drosophila*, one NURF55 homolog may function as a common platform for the assembly of protein complexes involved in chromatin metabolism (Martínez-Balbás et al. 1998). On the other hand, although several NURF55 homologs may be present, not all of them are involved in PRC2 function, as can be exemplified by the MULTICOPY SUPPRESSOR OF IRA (MSI) 1 – MSI5, the five NURF55 homologs in *Arabidopsis*, where only MSI1 has been associated with PRC2 function (Hennig et al. 2003; Köhler et al. 2003; Hennig et al. 2005). In summary, although we detect the general presence of NURF55 homologs throughout the eukaryotic lineages, future experimental work will be needed to associate the homologs with PRC2 function.

For this reason, several previous studies avoided computing NURF55 phylogenetic tree (Shaver et al. 2010; Zhao et al. 2020), to bypass confusing interpretations, and only E(z), ESC, and Su(z)12 subunit were used to infer the phylogeny of the PRC2 complex. Other studies include the NURF55 phylogenetic tree but within one organism or lineage (Huang et al. 2017; Juan et al. 2017). Here, we reconstruct the phylogeny of NURF55 in all eukaryotic lineages, using sequences identified through the PcG-finder and previously identified NURF55 sequences in KOG0264. Together, 830 sequences were obtained and used to construct a ML phylogenetic tree. The clustering of sequences in our tree (fig. S2C, Supplementary Material online) is distinct from the species tree of eukaryotes (Adl et al. 2012; Burki et al. 2020). A chaotic distribution of species in the tree can be explained by numerous putative Horizontal Gene Transfer (HGT) events (supplementary fig. S2C), which provide a transient advantage to the gene acceptors (Keeling and Palmer 2008). In addition, a separation of *A. thaliana* and *H. sapiens* NURF55 homologs was observed. Based on the location of these separated sequences in the tree, we can divide the tree into two groups (fig. S2C, Supplementary Material online). The first group of sequences clustered in the crown of the tree and includes the only *D. melanogaster* homolog (NP_524354.1), the three *A. thaliana* homologs (*MSI1* “AT5G58230”, *MSI2* “AT2G16780” and *MSI3* “AT4G35050”), and *H. sapiens* homologs (*RBBP7* “NP_001185648.1” and *RBBP4* “NP_001128727.1”). A similar separation between the NURF55 homologs has been observed before in plants (Huang et al. 2017; Ni et al. 2019). The second group of sequences clustered near to the root of the tree, including the rest of *A. thaliana* homologs (*MSI4* “AT2G19520.1” and *MSI5* “AT4G29730.1”) and newly identified *H. sapiens* homologs (*WDR73* “NP_116245.2” and *WDR59* “NP_085058.3”) (fig. S2C, Supplementary Material online).

We identified up to four WD-40 repeat domains in the NURF55 proteins (table S1F, Supplementary Material online) similar to the ESC proteins. Contrary to the ESC subunit, NURF55 homologs also include a CAF1C_H4-bd domain that is known to bind histone H4 to promote histone deposition onto newly synthesized DNA within CAF1 (Murzina et al. 2008; Ni et al. 2019) (table S1F, Supplementary Material online). Interestingly, both *D. melanogaster* and *H. sapiens* homologs clustered in the crown of the tree contain CAF1C_H4-bd domain while the rest of the *H. sapiens* homologs clustering near to the root of the tree do not contain this domain (table S1F; fig. S2C, Supplementary Material online). In contrast, all *A. thaliana* homologs contain the CAF1C_H4-bd domain even if they are phylogenetically separated, indicating that the presence of the CAF1C_H4 bd domain is not a determinant of the phylogeny position. Thus, the presence or absence of the CAF1C_H4-bd domain does not allow to distinguish between the NURF55 engagement in the CAF1 and/or the PRC2 complexes. Whether the phylogenetic position could distinguish homologs with specific chromatin complex engagement needs to be determined in the future. Similarly, it will be interesting to determine the potential redundancy of the *A. thaliana* MSI homologs in chromatin complexes (table S1F; fig. S2C, Supplementary Material online).

For better interpretation of this protein evolution, we compute an additional tree combing prokaryote sequences from several bacterial, archaeal, and Asgard-archaeal groups (fig. S2D, Supplementary Material online). The prokaryotic sequences were identified using PcG-finder with excluding the orthologous assignments step since the NURF55 protein does not assign to any bacterial Orthologous Group (OG). The rooted and well-supported tree shows a complete separation between the prokaryotic and eukaryotic domains with few exceptions (fig. S2D, Supplementary Material online). Only some homologs of Alveolata, Discoba, Stramenopoila, Cryprophyta, and Rhodelphea have retained the bacterial origin by clustering within the bacterial domain while few homologs of the Alveolata, Discoba, Stramenopoila, Viridiplantae, and Glaucophyta retain the cyanobacterial (plastid) origin (fig. S2D, Supplementary Material online). Even though many of the eukaryotic transfers in this tree remain unclear, the tree shows interesting gene transfer events such as the branching of the Archaeplastida (Rhodophyta, Rhodelphea, Viridiplantae, Glaucophyta, and Cryptophya) and SAR group sequences together near to the eukaryotic root of the tree (supplementary fig. S2D). More gene transfer events are found at the branching of Rhodophyta and Rhodelphea sequences with ciliates and the euglenids *Eutreptiella gymnastica* sequences near to Rhizaria and Cryprophyta sequences at the crown of the tree (fig. S2D, Supplementary Material online). This indicates that some homologs of this protein have been transferred multiple times through the complex endosymbiosis events (Keeling 2010; Qiu et al. 2013; Oborník 2019; Ponce-Toledo et al. 2019).

In summary, we identify homologs of PRC2 core subunits, and particularly E(z) homologs, in all eukaryotic lineages, including Discoba - a lineage currently placed proximal to the root of eukaryotes. This indicates early origin of the PRC2 complex or even presence in the LECA. Interestingly, we did not identify Su(z)12 homologs in Discoba and we find a larger number of species where E(z) and Esc homologs are identified than those where E(z) and Su(z)12 are identified (fig. S1B, Supplementary Material online). This suggests that E(z)-Esc may represent an ancient primary functional core module of the PRC2. Su(z)12 may have emerged later and its secondary loss seems more frequent than loss of ESC, suggesting its dispensability. This is in line with identification of E(z) and Esc homologs in all studied species in the green lineage, which suggested conservation of an ancient PRC2 composition (Huang et al. 2017). It must be noted however that the function of Su(z)12 homologs in vivo may be fulfilled by unrelated proteins, as is the case of the MES-3 subunit of the *Coenorhabditis elegans* PRC2 complex (Xu et al. 2001; Bender et al. 2004). Although the NURF55 homologs are well conserved in eukaryotes, their involvement in multiple chromatin-modifying complexes impedes concluding on their conserved participation in PRC2, and this aspect will require further study.

### Evolution of the E(z) homologs among the SET-domain proteins

E(z) is a part of the SET-domain protein family, which consists of seven subfamilies [PRDM, SMYD, SETD, SUV, ASH, Trx, and E(z)] and other unclassified SET proteins (Zhang and Ma 2012). SET-domain proteins display histone methyltransferase (HMTase) activities. Each targets specific lysine residues of histone H3 or H4 (Rea et al. 2000; Zhang and Ma 2012). To date, SET-domain proteins have been identified in bacteria and archaea (Alvarez-Venegas, 2014; Alvarez-Venegas et al., 2007; Aravind & Isyer, 2003; Stephens et al., 1998), but their presence in the Asgard group has not been investigated thus far. Phylogenetic analyses have shown that bacterial SET-domain homologs evolved independently of their eukaryotic counterparts (Alvarez-Venegas 2014); HGT of SET homologs between bacteria and eukaryotes were rejected (Alvarez-Venegas et al. 2007; Murata et al. 2007). Nevertheless, lateral gene transfer events of SET-domain homologs have been reported between bacteria and archaea (Alvarez-Venegas et al. 2007; Alvarez-Venegas 2014). We, therefore, asked how the identified E(z) homologs evolved in relation to other representatives of SET-domain protein families.

To explore the eukaryotic and prokaryotic SET-domain protein evolutionary relationships, we traced back the SET-domain proteins in prokaryotes. For this analysis, we used a modified PcG-finder. The hits mapping to the bacterial Orthologous Group (OG) cluster of the SET-domain protein (COG2940) were selected in the orthologs assignment step. The SET-domain proteins were identified in 41 out of 105 bacterial species studied (39%), representing 26 different taxonomic groups (table S2C, Supplementary Material online). We found the SET-domain proteins only in the Euryarchaeota of all the archaeal groups (table S2C, Supplementary Material online). For the first time, we identified a SET-domain protein in one of the Asgard species (*Candidatus Lokiarchaeota archaeon*) (table S2C, Supplementary Material online). Our results show that the SET-domain proteins are less conserved in Bacteria and Archaea than what was reported previously (Alvarez-Venegas 2014), likely due to a lower number of representative species per group in our analysis.

The comprehensive ML phylogenetic tree was computed based on all the E(z) sequences (supplementary table S1C) and prokaryotic SET-domain protein sequences (table S2C, Supplementary Material online) identified in this study and other sequences of proteins from different SET-domain protein subfamilies identified previously in *Arabidopsis thaliana, Drosophila melanogaster* and *Homo sapiens* (Dillon et al. 2005; Ng et al. 2007; Zhang and Ma 2012; Sarma and Lodha 2017) (fig. 3). The tree was rooted using the Asgard homolog (TFG06932_1), to identify the evolutionary direction of the proteins (fig. 3; table S2C, Supplementary Material online).

**Figure 3.**
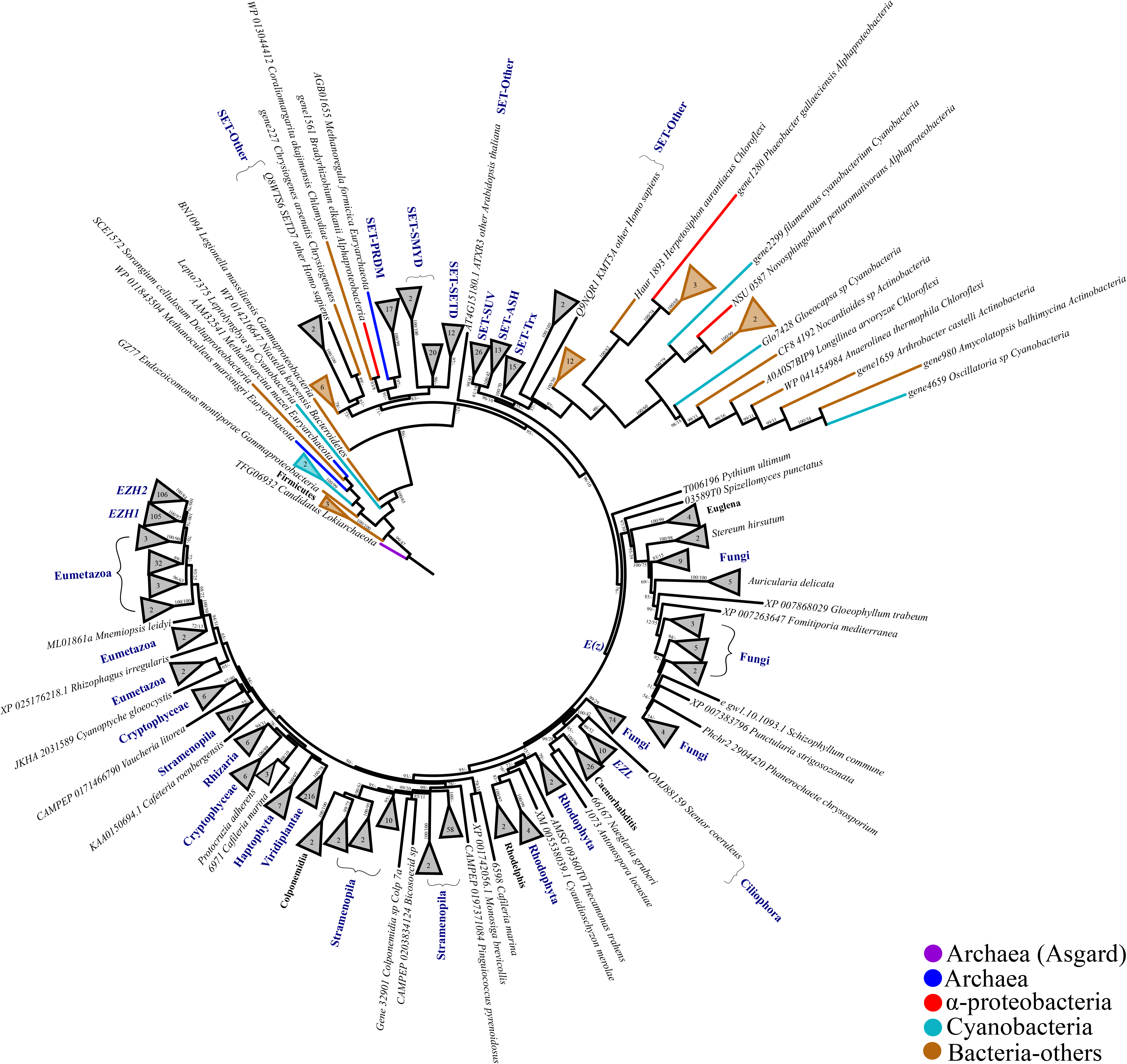
A ML phylogenetic tree based whole protein sequence, showing the evolutionary relationships of prokaryotic and different eukaryotic sub-families of SET-domain proteins. The tree was rooted with the Asgard SET-domain protein and the maximum likelihood branch support values are given in % (IQ-TREE/RAxML-NG).

In contrast to the conclusions of previous work (Alvarez-Venegas 2014), we detected three HGT events (fig. 3). Interestingly, two of them include the unclassified SET-domain sequences. Our analysis shows that this class of proteins (other SET-domain proteins) has a bacterial origin, suggested by the de-clustering of these proteins with any of the other SET-domain subfamilies in the previous studies (Dillon et al. 2005; Ng et al. 2007; Zhang and Ma 2012). The third HGT was between the mammalian SET-domain subfamily (PRDM) (Vervoort et al. 2016) and the euryarchaeota *Methanoregula formicica* homolog (AGB01655.1); forming a sister group with alphaproteobacteria (1038859.AXAU01000009_gene1561) and *Chlamydiae* (WP_013044412_1) homologs (fig. 3; table S2C, Supplementary Material online). A HGT event between eukaryotic and *Chlamydiae* SET-domain proteins has been identified previously (Stephens et al. 1998). Moreover, a recent report highlighted the contribution of *Chlamydiae* on the evolution of eukaryotes (Stairs et al. 2020). In line with previous analyses, our phylogenetic tree identified lateral gene transfer events between archaea and bacterial sequences (Alvarez-Venegas 2014) as well as transfer events between different bacterial groups (fig. 3). Importantly, the sequences of SET-domain proteins within each subfamily formed monophyletic groups without mixing (fig. 3) as in previous phylogenetic analyses (Huang et al. 2017; Chen et al. 2020), and the relationships and topology of the SET-domain subfamilies were highly similar. Interestingly, the E(z) subfamily clustered in the crown of the tree without interaction with the prokaryotic sequences and shows the same topology as the eukaryotic-E(z) tree (fig. 2). The branch containing E(z) homologs forms a well-separated sister branch to the SET-domain subfamilies SET-SUV, SET-ASH and SET-Trx; and all these sub-families branch separately from the SET-PRDM, SET-SMYD and SET-SETD sub-families. The E(z) subfamily of SET-domain proteins therefore seems to have diverged following the emergence but preceding the expansion of eukaryotes, providing a suitable model to address the evolution of PRC2 catalytic function.

We next addressed the question of whether differences in the sequences of the SET-domains alone follow the same phylogeny and separation of E(z) homologs into 5 clades; and whether this may potentially indicate functional separation (e.g. substrate specificity) of the SET-domain proteins. To this end, we extracted the amino acid sequences comprising the SET-domain using an in-house python script (SET extractor). Next, a rooted-ML tree was computed using the extracted SET-domain sequences (supplementary fig. S3) from the same dataset as was used to compute the full-length SET-domain proteins phylogeny (fig. 3). Both phylogenetic trees show similar topology; and in particular the clustering of the E(z) subfamily members into five clades is retained. Still, we identified a few interesting topology differences in the reconstructed phylogeny of the SET-domains alone. For instance, the glaucophyte *Cyanoptyche gloeocystis* domain-sequence was clustered in the root of all E(z)-SET-domain sequences, but this branching is not supported as many tree nodes (supplementary fig. S3); this might be due to the short length of the domain sequences. Moreover, the sequence of the alphaproteobacteria *Pelagibacter ubique* clustered with the rhodophyte *Cyanidioschyzon merolae* within the clade III that includes all other rhodophytes. This could indicate that in clade III of E(z) homologs, the SET domain retained bacterial features. Finally, some SMYD-SET domain sequences branched on the root of clade V of E(z) SET-domains (fig. S3, Supplementary Material online). This divergence might be a consequence of the contribution of the SMYD-SET domain subfamily to histone modifications that can be both repressive and activating (Tracy et al. 2018).

SET-domains display different substrate specificity (Herz et al. 2013). Interestingly, this also holds true for different SET-domains within the E(z) subfamily, as some E(z) homologs have been demonstrated to methylate substrates other than H3K27 (Frapporti et al. 2019), and they can work as mono-/di-methyltransferases in mammals (Lee et al. 2013) or solely as tri-methyltransferases in plants (Wang et al. 2021). We therefore asked whether the clustering of the SET-domains and the separation of the E(z) SET-domains into the different clades may correlate with substrate specificity. The SET-domain consists of a series of conserved sequence patterns (motifs) such as GxG, YxG, RFINHxCxPN, ELxFDY, and the last two motifs form a pseudo-knot-like conformation in the fold that has specific roles in binding and catalysis (Cheng et al. 2005) (fig. 4). In the C-terminal area, this pseudoknot conformation brings together the sequence motifs RFINHxCxPN and ELxFDY, forming the active site close to the unique cofactor binding pocket. This binding pocket is crucial for the specific interaction of the methyl donor (cofactor) with H3 at specific locations through a narrow hydrophobic deep channel, which is connected to the peptide-binding cleft, located exactly at the opposite surface of the SET-domain (Dillon et al. 2005).

**Figure 4.**
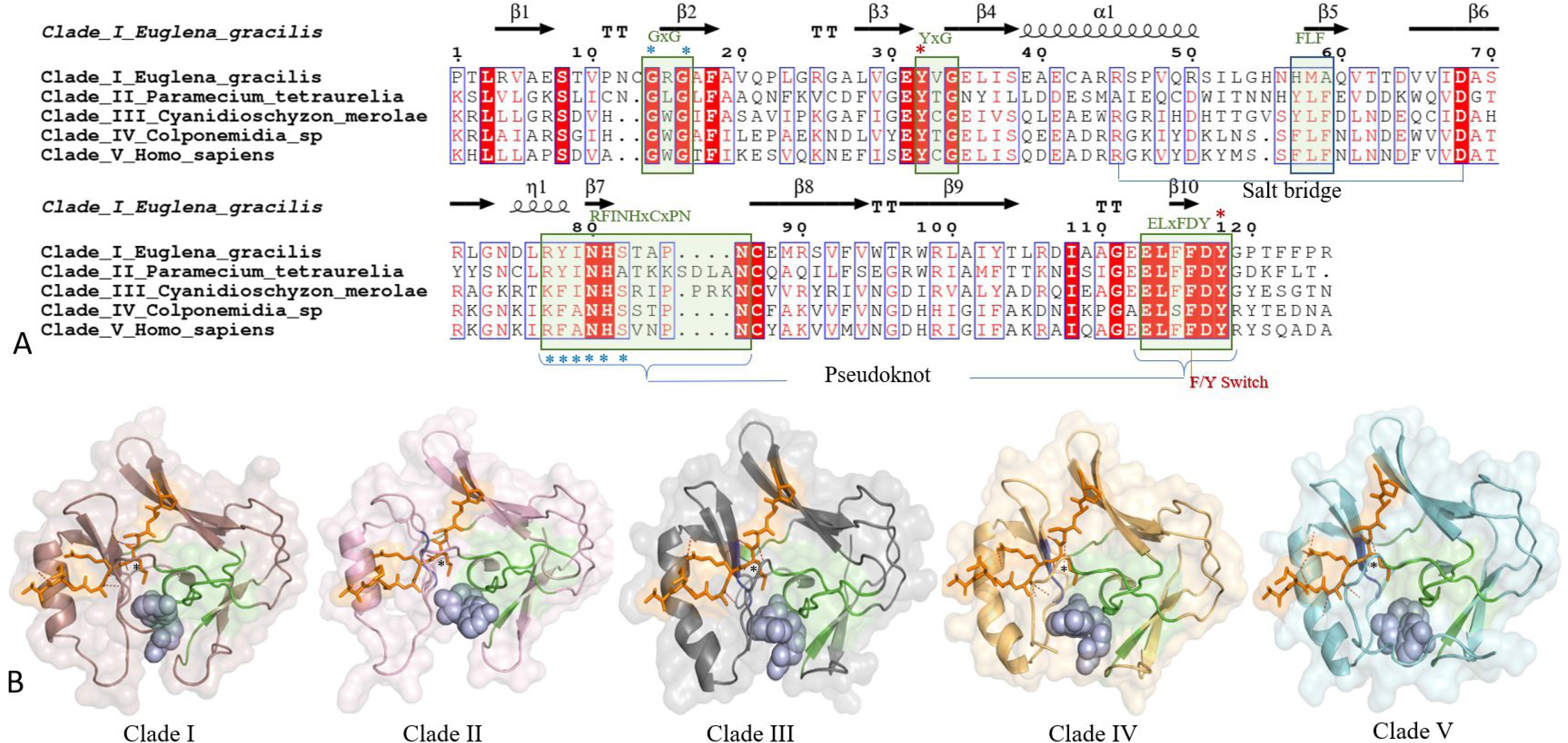
Sequence-structure comparison of representative SET-domains from five E(z) clades **A)** Alignment of SET domains of different clade representatives. Invariant (absolutely conserved) residues are highlighted in red background with white text. Conserved residues are highlighted in red font. Conserved signature motifs I-IV (Green box) and FLF (Blue box), salt bridge that causes intramolecular interaction, pseudoknot, F/Y switch controlling methylation (mono-, di- or tri-methylation), catalytic sites (red asterisk), cofactor binding sites (blue asterisk) are highlighted. **B)** Cartoon/surface view of the SET domain of each clade. Cofactor (light blue) and the substrate analog (H3K27M-peptide inhibitor; orange color) are represented by stick and sphere form. The position of ‘K27M’ and lysine binding channel is indicated by a black asterisk. Peptide binding cleft area highlighted in light orange. Conserved i-IV signature motifs are highlighted in green and the FLF motif in blue. Interactions of domain with peptides (hydrogen bonds) are represented by red dashes).

The catalytic SET-domain sequences (120-130 amino acids) are substantially conserved within individual subfamilies and E(z) clades (fig. S4, Supplementary Material online). Among the different SET-domain subfamilies, SET-domains of all E(z) clades are well conserved with identical folds and similar binding pattern. Besides, SET-domains of TRX and ASH protein subfamilies (~120 amino acid) are also adopting almost similar structural fold as that of E(z)-SET domain, supporting their phylogeny-structure relationship. Meanwhile, the SET-domains of PRDM, SETD, SMYD, and SUV subfamilies are quite distinct in terms of sequence length, size and structure. These results are in line with the SET-domain phylogenetic tree (fig. S3, Supplementary Material online), which show that the SET-domain sequences of PRDM, SETD and SMYD sub-families are more divergent within the tree with respect to the E(z) subfamily. Furthermore, these findings support the functional divergence of PRDM and SMYD sub-families (Hohenauer and Moore 2012; Vervoort et al. 2016; Tracy et al. 2018) and highlight a possible functional divergence of the SETD subfamily. Our phylogenetic analyses and sequence conservation pattern clustered the E(z) SET-domain sub-family into five clades (fig. 3 and 4; fig. S3 and S4, Supplementary Material online). Among these clades, the experimentally characterized SET-domain of human E(z) (Justin et al. 2016) is located in clade V. Further, the SET-domains of all E(z) clades (clade I-V) exhibit significant sequence/structural similarities with human EZH2-SET-domain. Some sequence conservation patterns were identified in each of the E(z) clades, which may contribute to the differentiation of clades within the E(z) sub-family (fig. 4A; fig. S4, Supplementary Material online). Generally, four sequence motifs (GxG, YxG, RFINHxCxPN, and ELxFDY) are highly conserved in all E(z) SET-domains while some minute alterations were observed among the five E(z) clades (fig. 4A; supplementary fig. S4). Two conserved motifs (GxG and YxG) are present in the N-terminal area and proved to have significant roles in catalysis. GxG motif is located at the end of the first β-strand and conserved in all E(z) SET-domain clades (fig. 4A; fig. S4, Supplementary Material online). This motif is shown to have significant roles in the interaction with the methyl donor (Joshi et al. 2008). YxG motif is a catalytic motif that also showed a high conservation pattern with some alterations in the position of Glycine (G→A or G→S or G→E) (fig. 4A; supplementary fig. S4) that may cause a change in conformation and/or interaction. Towards the C-terminal end the RFINHxCxPN, and ELxFDY motifs show many alterations in each clade. Finally, the hydrophobic motif “FLF”, that contributes to the lysine binding pocket (Joshi et al. 2008), is conserved in clades III and V while this motif is lost in many of the Clade I and II sequences. All the above-mentioned motifs are highly conserved in clade V that contains E(z) SET domains of higher eukaryotes, as compared to other E(z) clades that mainly contain SET-domains of lower eukaryotes (fig. 4A; fig. S4, Supplementary Material online).

Even though E(z) clades show alterations in amino acids, they essentially adopt a similar structural fold and cofactor binding, with minor/or no alterations in the peptide-binding cleft, which may contribute to the methylation pattern (fig. 4A; fig. S5, Supplementary Material online). Modeled structures of representatives of each of these clades show an identical three-dimensional structural fold with a conserved anti-parallel β-barrel and slightly variable cofactor binding pocket that comprises the enzyme active site residues (fig. 4B). During catalysis by the evolutionary conserved SET-domain (fig. S5A, Supplementary Material online), the methyl group is transferred from the cofactor (S-adenosylhomocysteine (SAH)) to the substrate peptides through the hydrophobic deep lysine channel (Dillon et al. 2005). We next asked whether the peptide binding and cofactor interaction in the catalytic SET-domain could define the SET-domains of the five E(z) clades and may contribute to the differences in substrate specificity. We modeled the structures of the different E(z) SET-domain clades, either in the apo-(no ligand) or holo-form when bound to the cofactor S-adenosyl homocysteine (SAH) and a peptide substrate analog (H3K27M), mimicking the domain/substrate/cofactor complex (fig. 4B; supplementary fig. S5A). The above-mentioned RFINHxCxPN, and ELxFDY motifs in all the E(z) SET-domain clades form a specific unusual knot-like (‘pseudo-knot’) structure, which contains the specific short NHS-motif through which it interacts with the cofactor via hydrogen bonding. Nevertheless, the cofactor (SAH) and substrate/inhibitor bind at different clefts on the opposite surfaces of the SET-domain (fig. S5B, Supplementary Material online). Generally, the interaction pattern of all E(z)-domain representatives excluding clade II showed a similar binding pattern for both cofactor and peptide (fig. 4B; fig. S5A, Supplementary Material online). Interestingly, a portion of the inhibitor (M of H3K27M, also K in H3K27Me) anchors deeply into the hydrophobic lysine channel, towards the direction of the cofactor binding pocket, while other residues are mostly exposed (fig. 4B; fig. S5A, Supplementary Material online). Selectivity/methylation pattern is also based on the residues present in the lysine channel (fig. S5A, Supplementary Material online). In the modeled structure of clade II, some of the loops/turns near the cofactor binding pocket are slightly disordered when compared to other models, which might be the reason for the binding difference (fig. S5A, Supplementary Material online).

Surprisingly, the structure model of the glaucophyte *Cyanoptyche gloeocystis* SET-domain shows significantly different anchoring patterns for H3 peptide substrate, and deep anchoring for ‘K27M’ into the cavity was not observed, even though the structural fold showed substantial identity to the other E(z)-SET domains (fig. S6, Supplementary Material online). Interestingly, the structure model shows better interaction with H3K9M and ‘K9M’ was directed towards the channel which suggests the putative possibility of H3K9me catalysis (fig. S6, Supplementary Material online). Previously, H3K9me (as well as H3K27me) was experimentally shown to be carried out by the Ezl homolog of *Paramecium tetraurelia* (clade II). Interestingly, the modelled orientation of both the H3K9M and H3K27M in the *P. tetraurelia* SET-domain suggest methylation events. These findings are in line with the respective phylogenetic positions of these SET-domains (at the root of the E(z) SET-domain sub-family in *C. gloeocystis* and early-diverged clade II for *P. tetraurelia* - fig. S3 and S6, Supplementary Material online), raising indications on possible functional evolution of the E(z) SET-domains.

## Conclusion

Despite the intensive investigation of PRC2 core subunits distribution in plants and opisthokonts, the diversity of these subunits within early branching eukaryotes remains enigmatic. Here, we present the first phylogenetic profiling of PRC2 core-subunits across all eukaryotic lineages using a computational automated tool (PcG-finder) which can be used in future studies. Altogether, our results strongly suggest that the homologs of the catalytic subunit E(z) and the core subunits ESC and NURF55 existed in lineages hypothetically closest to the eukaryotic root (fig. 5), in agreement with the standing hypothesis of the presence of PRC2 in the LECA. We identify the co-occurrence of the homologs of E(z) and ESC more frequently than E(z) and Su(z)12 and fail to identify Su(z)12 homologs in Discoba and Metamonada, proposing that the E(z)-ESC module may have been the initial PRC2 functional module. NURF55 is the most conserved subunit within all eukaryotic lineages, supporting its involvement in other chromatin-related complexes. Su(z)12 is the most divergent subunit (fig. 5), indicating its potential dispensability or functional substitution. Moreover, we could not associate the loss of any of the subunits either with the uni- or multi-cellularity or lifestyle of species within the studied eukaryotic lineages (fig. 5). However, our findings highlight important future questions that need to be addressed experimentally, including PRC2 catalytic activity, E(z) substrate specificity and biological significance of the PRC2. Additionally, future work with more advanced versions of genome assemblies will be needed to settle the question of secondary losses of E(z) in individual species of different lineages or even major eukaryotic lineages such as the alveolates (fig. 5).

**Figure 5.**
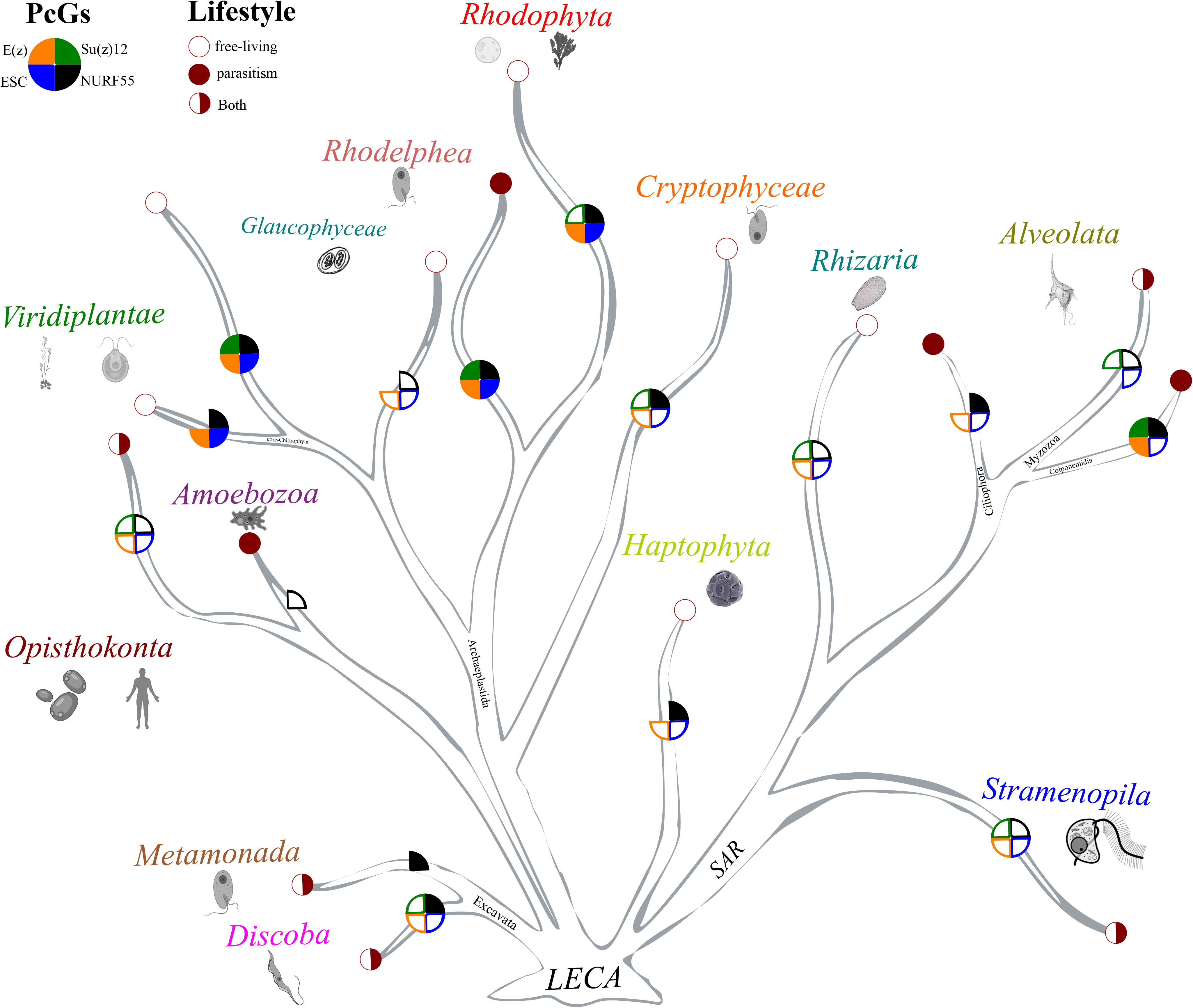
Schematic of the eukaryotic tree of life shows the summary of PRC2 subunit diversity.

We present a robust phylogenetic analysis that includes all SET-domain proteins in prokaryotes, all E(z)-subfamily SET-domain proteins and representative proteins of other SET-domain protein families in eukaryotes. The detection of SET-domain protein homologs in Asgard species will allow a better understanding of the mechanisms involved in lysine methylation in prokaryotes and will clarify the evolutionary relationships among methyltransferases from the four domains of life. Both full-length and SET-domain sequence-based phylogenetic reconstruction highlighted the existence of five clades of E(z)-SET-domain proteins. While there are amino acid sequence signatures of each of the SET-domain clades, representative SET-domains of all clades adopt a similar structural fold and display interaction potential for the substrate H3K27M. Interestingly however, SET-domains located proximal to the root of the E(z)-SET domains display efficient interaction with H3K9M, that may indicate potential differences in substrate specificity and its evolution.

## Materials and Methods

### Data sources

The complete predicted proteome sequences of studies organisms (table S1A, Supplementary Material online) were obtained from JGI (http://genome.jgi.doe.gov), the Eukaryotic Pathogen database (https://eupathdb.org) and NCBI GeneBank (https://ncbi.nlm.nih.gov/). The Marine Microbial Eukaryote Transcriptome Sequencing Project database (MMETSP) (Keeling et al. 2014) and the 1000 Plants (1KP) (https://sites.google.com/a/ualberta.ca/onekp/) (Leebens-Mack et al. 2019) were additional sources for predicted proteome sequences inferred from transcriptomic data. Some proteome sequences were retrieved using mining publications (*Rhodelphis marinus, Rhodelphis limneticus, Euglena gracilis var. Bacillaris* and metamonads from Leger et al. (Leger et al. 2017) or kindly made available by our collaborators. In proteome datasets, when two or more protein sequences at the same locus were identical and overlapping, the longest sequence was considered.

### PcG-finder pipeline

The pipeline we established and called “PcG-finder” is composed of three different steps: Homology search, orthology assignments and domains architecture scanning (fig. 6). The whole pipeline was implemented in single python program and deposited in github.

**Figure 6.**
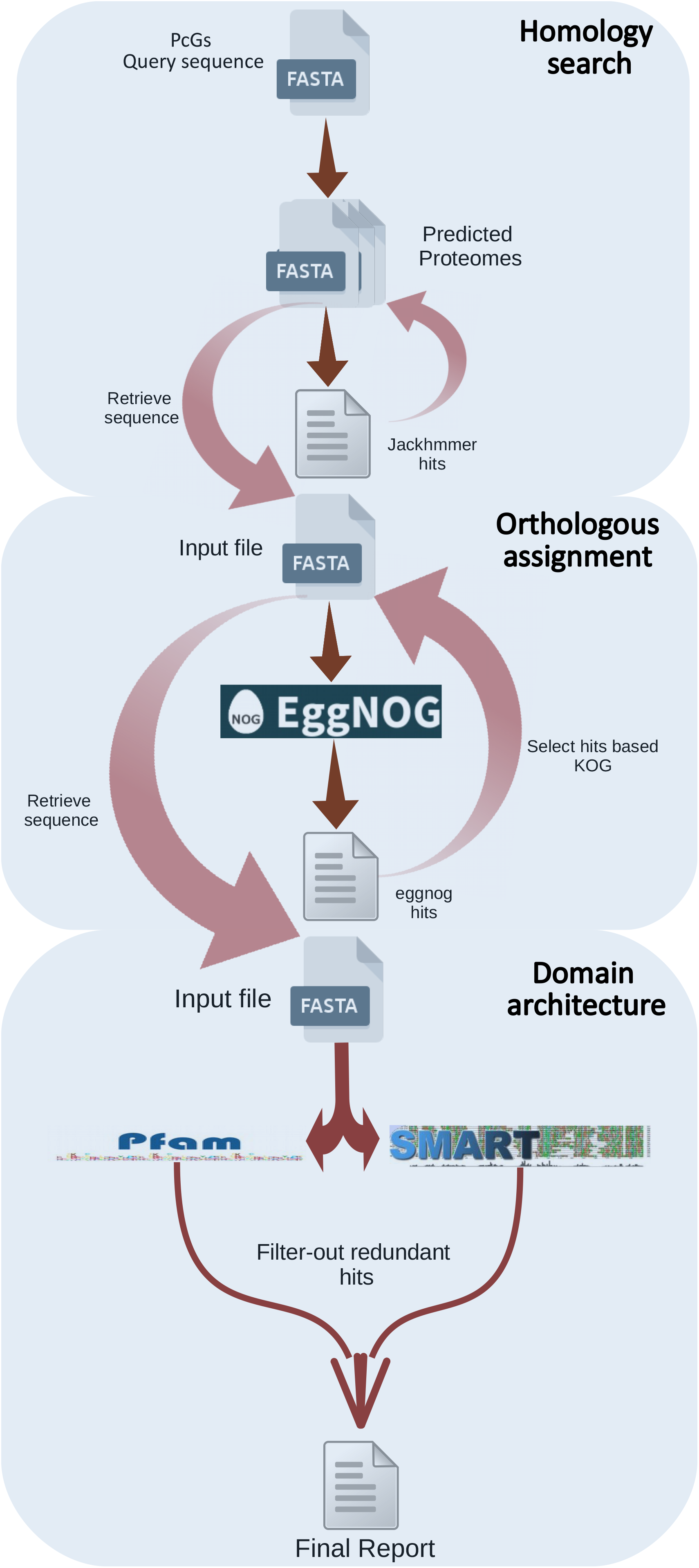
Schematic flow chart of the PRC2 subunits prediction pipeline (PcG-finder).

The *Drosophila* E(z) (NP_001137932.1) reference amino acid sequence was used to search of all predicted proteome sequences retrieved previously using the Hidden Markov model (HMM)-based tool jackhammer (Johnson et al. 2010). The highest scoring protein from each target organism was then used for a jackhmmer search against the reference genomes of *Arabidopsis thaliana, Drosophila melanogaster* and *Homo sapiens*, to confirm a reciprocal best match to the original protein. For the other members of PRC2, jackhmmer searches were performed using *Drosophila* reference amino acid sequences Su(z)12 (NP_730465.1), ESC (NP_477431.1) and NURF55 (NP_524354.1) (fig. 6).

Evolutionary genealogy of genes: Non-supervised Orthologous Groups (eggNOG) mapper was used for hierarchical resolution of orthology assignments and fine-grained relationships based on phylogenetic analysis. eggNOG mapper implements a species-aware clustering algorithm based on the concept of triangulation of best reciprocal hits (Tatusov et al. 1997) is applied to identify Orthologous Groups (OGs): sets of homologous sequences that started diverging from the same speciation event. In addition, manually curated OGs available for the eukaryotes (KOGs) (Bioinformatics et al. 2003) were integrated into the corresponding levels in eggNOG. Only eggNOG-hits with KOG1079, KOG2350, KOG1034 and KOG0264 for E(z), ESC, Su(z)12 and NURF55 respectively were selected. These KOGs were identified earlier using *Drosophila’s* reference amino acid sequences (fig. 6).

Finally, the SMART and Pfam databases were employed to identify conserved domains present in E(z), ESC, Su(z)12 and NURF55 from different organisms (Letunic and Bork 2018; El-Gebali et al. 2019), both SMART and Pfam databases were merged, and redundant domains were filtered-out and the Hidden Markov model (HMM)-based tool hmmscan (https://github.com/EddyRivasLab/hmmer) was used to scan domains architecture. Only sequences with the catalytic or conserved domain of the references were retained.

### Phylogenetic analyses

Both KOG sequences and PcG-finder true positives sequences were aligned using MAFFT software (Katoh and Standley 2013) and ambiguously aligned regions were excluded from further analysis using trimAl software (Capella-Gutiérrez et al. 2009). Alignments were tested using ProtTest v3 (Darriba et al. 2011) to choose the appropriate model for amino acid substitution.

Two separated Maximum likelihood (ML) phylogenetic trees were computed using RAxML-NG (Kozlov et al. 2019) and IQ-TREE 2 (Minh et al. 2020) software. ML analyses were performed using 1000 bootstrap replicates. The supporting values from both software were noted on the ML-unrooted tree. Only SET-protein phylogenetic trees were rooted with Asgard sequence, which was considered as the out-group.

### Sequence-structure analysis

The representative sequences of each subfamily or clade of SET-domain were aligned independently using MAFFT software (Katoh et al. 2019) based on the L-INS-I method. ESpript v. 3.0 (Robert and Gouet 2014) was used to visualize the sequence conservation pattern. To get insights into the structural features, protein models based on homology were generated for representative species by i-Tasser (Roy et al. 2010). The model quality was confirmed using PROCHECK (Laskowski et al. 2012). Further, these structurally validated homology models, as well as experimentally characterized models, were used to generate sequence-structural alignments using the ENDscript v. 2.0 (Robert and Gouet 2014). In silico interaction using the peptide inhibitor (H3K27M or H3K9M) and methyl donor (SAH-cofactor) were predicted based on the experimentally characterized human EZH2 SET-domain (PDB ID: 5HYN). H3K27Me was also used as a comparative analysis. SwissDock (Grosdidier et al. 2011) and HawkDock server (Weng et al. 2019) were used for docking and Pymol v. 2.4.1 (https://pymol.org/2/) and Molsoft ICM-Browser (Neves et al. 2012) were used for molecular visualization and comparative analysis between the modeled and reference protein structures.

## Acknowledgments

This work was supported by the Czech Academy of Sciences, ERC-CZ, grant number [ERC200961901] to IM. Computational resources were provided by the CESNET LM2015042 and the CERIT Scientific Cloud LM2015085, funded under the programme “Projects of Large Research, Development, and Innovations Infrastructures”. The authors thank Dr. Aleš Horák for providing the transcriptomic data for the some unidentified diplonema and stramenopile species.

## Author Contributions

AS, IM and MO conceived the research. IM acquired funding. AS and MV performed the bioinformatics analyses. AS, IM and MV drafted the manuscript. All authors revised the first draft and read and approved the final version of the manuscript.

## Data Availability

PcG-finder pipeline python-software available at https://github.com/Iva-Mozgova-Lab/PcG_finder and SET-extractor script available at https://github.com/Iva-Mozgova-Lab/SET_extractor.

## Notes

### Competing Interest Statement

The authors have declared no competing interest.

